# Sub-Millimetre Resolution Laminar Fmri Using Arterial Spin Labelling in Humans at 7 T

**DOI:** 10.1101/2020.08.22.261693

**Authors:** Sriranga Kashyap, Dimo Ivanov, Martin Havlicek, Laurentius Huber, Benedikt A. Poser, Kâmil Uludağ

## Abstract

Laminar fMRI at ultra-high magnetic field strength is typically carried out using the Blood Oxygenation Level-Dependent (BOLD) contrast. Despite its unrivalled sensitivity to detecting activation, the BOLD contrast is limited in its spatial specificity due to signals stemming from intra-cortical ascending and pial veins. Alternatively, regional changes in perfusion (i.e., cerebral blood flow through tissue) are colocalised to neuronal activation, which can be non-invasively measured using arterial spin labelling (ASL) MRI. In addition, ASL provides a quantitative marker of neuronal activation in terms of perfusion signal, which is simultaneously acquired along with the BOLD signal. However, ASL for laminar imaging is challenging due to the lower SNR of the perfusion signal and higher RF power deposition i.e., specific absorption rate (SAR) of ASL sequences. In the present study, we present for the first time in humans, isotropic sub-millimetre spatial resolution functional perfusion images using Flow-sensitive Alternating Inversion Recovery (FAIR) ASL with a 3D-EPI readout at 7T. We show that robust statistical activation maps can be obtained with perfusion-weighting in a single session. We observed the characteristic BOLD amplitude increase towards the superficial laminae, and, in apparent discrepancy, the relative perfusion profile shows a decrease of the amplitude and the absolute perfusion profile a much smaller increase towards the cortical surface. Considering the draining vein effect on the BOLD signal using model-based spatial ‘convolution’, we show that the empirically measured perfusion and BOLD profiles are, in fact, consistent with each other. This study demonstrates that laminar perfusion fMRI in humans is feasible at 7T and that caution must be exercised when interpreting BOLD signal laminar profiles as direct representation of the cortical distribution of neuronal activity.

## INTRODUCTION

Neuronal activity in the brain is associated with an increased metabolic demand accompanied by changes in haemodynamics such as blood oxygenation, flow and volume (for reviews see: [1–4]). Functional magnetic resonance imaging (fMRI) is a technique that can non-invasively measure these changes and allows inferring the spatial pattern of neuronal activity while performing a task or at rest. Improvements in MRI technology over the past decades, such as higher magnetic field strengths, novel sequences, optimised pulse designs, and parallel imaging, have pushed the spatial and temporal limits to an extent wherein MRI at ultra-high magnetic field (UHF, ≥7T) can routinely achieve sub-millimetre spatial resolution voxels in humans, for both structural and functional imaging (see Special Issues: [5,6] and reviews therein). While fMRI investigations have yielded robust, reproducible functional parcellation [7] of different brain areas consistent with previous ex vivo cyto- and myelo-architectural studies [8,9], the advantages of UHF fMRI have enabled neuroscientists to investigate the mesoscopic circuitry within regions across cortical depths and, to a lesser extent, columns in humans (see Special Issue: [10] and reviews therein).

A vast majority of standard-resolution and laminar fMRI studies have been performed using the Blood Oxygenation Level-Dependent (BOLD) contrast [11,12]. While the BOLD contrast excels in its sensitivity to detect signal changes due to its high signal-to-noise (SNR), it is inherently limited in its spatial specificity relative to site of neuronal activation because of strong signal bias introduced via the intra-cortical ascending veins [13] and by the non-local signal spread (drainage effect) through pial veins [14,15]. Studies investigating the specificity of the laminar BOLD response in humans and animals [16–20] have consistently observed the largest signal change in the BOLD signal at the superficial layers and pial surface despite the fact that the peak of the neuronal activity is expected in the input layers (layer IV in human V1) for feed-forward stimuli [21,22]. Some earlier studies have investigated the leakage of the signal between laminae during steady-state [22–24]. Recently, a fully dynamical model of the laminar BOLD signal has been developed [13] that enables model-driven “deconvolution” (i.e. removal of the intra-cortical ascending venous signal) of the measured BOLD signal profiles to unravel the underlying neuronally-driven signal profiles. However, theoretical assumptions of these model-driven approaches have not yet been subjected to experimental validation.

The versatility of MRI provides the means to also measure other (non-BOLD) haemodynamic response parameters such as cerebral blood volume (CBV) using vascular space occupancy (VASO) [25–27] or cerebral blood flow (CBF) through tissue (perfusion) using arterial spin labelling (ASL) [28–30]. Most studies using these non-BOLD approaches have been carried out in animal models [3,4,31] and have only been applied to high-resolution human studies with the advent of UHF fMRI [32–34]. From the perspective of laminar fMRI, animal studies have shown that perfusion-weighting is a highly desirable contrast, even more so than total CBV, due to its spatial proximity to neuronal activation [18,35]. While CBV-weighted imaging using VASO has seen a resurgence for laminar fMRI applications [36], perfusion-weighted fMRI using ASL has been mostly limited to relatively low spatial resolution (≅ 2-4 mm) studies [37] (but see [38]). Achieving higher spatial resolutions, let alone sub-millimetre resolutions, with perfusion-weighting and adequate brain coverage is challenging. This is due to the relatively lower SNR of the perfusion-weighted signal owing to the low microvascular density and T_1_ recovery of the labelled arterial water signal, and the higher RF power deposition of ASL sequences in general. The SNR limitation can be addressed to some extent by moving to UHF. The gain in SNR due to increased field strength [39] and the prolonged longitudinal relaxation times (T_1_) [40,41] allows longer post-labelling delays, thereby, improving the perfusion SNR [33]. Recent developments using ASL at 7 T [32,33,42–44] have enabled pushing the spatial resolution for perfusion-mapping to the sub-millimetre regime [45,46] by overcoming several technical challenges; i.e. optimisation of sequence and pulse design [34,44,47], using dielectric pads [48] in order to improve the labelling efficiency [33], and utilisation of a 3D-EPI readout [49].

Taking together these advantages at UHF, the spatial specificity of the perfusion signal and the fact that ASL acquires both BOLD and perfusion-weighted images simultaneously makes ASL a very attractive tool for laminar fMRI. In the present study, we build on our previous work to acquire, for the first time, sub-millimetre resolution simultaneous BOLD and perfusion-weighted fMRI of the human visual cortex at 7T. We demonstrate that robust, participant-specific, single-session, high-resolution perfusion activation maps can be obtained for laminar fMRI in humans at 7T. We probe the cortical depth-dependence of BOLD and perfusion-weighted signals in response to visual stimulation in humans and reconcile our experimental findings using the recently proposed dynamic model of the laminar BOLD signal.

## METHODS

Seven healthy volunteers (median age=28 years) participated in the study following screening and having given written informed consent. The study was approved by the Ethics Review Committee for Psychology and Neuroscience (ERCPN) at Maastricht University and all procedures followed the principles expressed in the Declaration of Helsinki.

### Data acquisition

Data were acquired on a whole-body Siemens Magnetom 7T research scanner with a gradient system capable of maximum gradient amplitude of 70 mT/m and maximum slew rate of 200 T/m/s (Siemens Healthineers, Erlangen, Germany) and a 32-channel receive phased array head coil (Nova Medical, USA). The participant placement and preparatory procedure followed the protocol previously described in [33,42]. In short, the eye centres were taken as iso-centre reference (instead of the eyebrows, as is typically done) and supplementary cushions were provided to the participants under the neck, to ensure that the large feeding arteries to the brain were parallel to the B_0_. In addition, two 18×18×0.5 cm^3^ high-permittivity dielectric pads containing a 2.8:1 solution of calcium titanate (CaTiO_3_) and heavy water (D_2_O) by weight [50] were placed on either side of the head at the level of the participant's temporal lobes to increase B_1_ (therefore, labelling) efficiency at 7T [51].

#### Stimulus paradigm

Full contrast black-and-white radial flickering checkerboard was presented using PsychoPy v 1.90.0 [52] for 20 s (stimulus on) followed by 40 s of an iso-luminant grey background (stimulus off). Each functional run lasted ~12 minutes consisting of a 30 s initial baseline period and ten stimulus on-off blocks. The participants were instructed to remain motionless and fixate on a central fixation dot throughout each of the four functional runs.

#### Anatomical MRI

Anatomical data were acquired using a 3D-MP2RAGE [53] at 0.9 mm isotropic spatial resolution (192 sagittal slices; GRAPPA = 3; FoV_read_ = 230 mm; phase-encoding = A≫P; TI_1_/TI_2_ = 900/2750 ms; a_1_/a_2_ = 5°/3°; TE/TR = 2.39/4500 ms; partial-Fourier_phase_ = 6/8; bandwidth = 250 Hz/px; echo-spacing = 6.6 ms, TA = 6 min).

#### Functional MRI

Functional data were acquired at 0.9 mm isotropic resolution using a pulsed ASL (PASL) sequence [29] with a 3D-EPI readout [49] employing a FAIR [54] QUIPSS II [30] labelling scheme (44 axial slices; GRAPPA = 4; FoV_read_ = 192 mm; phase-encoding = A≫P; TE/TR = 15/2850 ms; a=19°; TI_1_/TI_2_ = 700/1891 ms; partial-Fourier_phase_ = 5/8; partial-Fourier_slice_ = 7/8; Ref. lines PE = 64; Ref. scan mode = FLASH [55]; bandwidth = 1124 Hz/px; echo-spacing = 1.02 ms, repetitions = 230, TA =11 min). The labelling was achieved using a tr-FOCI inversion pulse [47] (10ms) that provided efficient (up to 95%) slab-selective inversion despite inhomogeneous B_1_ and SAR constraints at high field [33,36]. Immediately after each of the four functional runs, five volumes with opposite phase-encoding were acquired for run-wise distortion-correction. All the ASL data were reconstructed using GRAPPA kernel of size {3,2} [42] and 8 iterations of the POCS algorithm [36,56]. The functional data acquisition slab was oriented to cover as much of the occipital lobe as possible in all participants centred on the calcarine sulcus (S3 Fig a).

### Data processing

The anatomical data were pre-processed in SPM12 r7487 (https://www.fil.ion.ucl.ac.uk/spm/software/spm12/) [57,58] and FSL v. 6.0 (https://fsl.fmrib.ox.ac.uk/fsl/fslwiki) [59,60]. This anatomical pre-processing workflow was developed particular to work well for MP2RAGE data. First, the second inversion image of the MP2RAGE was subjected to the automated segmentation in SPM12 [61]. The bias-corrected second inversion image was used to create a whole-brain mask using FSL BET [62]. The thresholded non-brain tissue classes from the SPM12 segmentation were summed together to create a mask of the non-brain tissue and large sinuses (for illustration of the analysis steps, see S1 Fig). The non-brain mask was manually curated in cases, in which the automatic masks were sub-optimal. The T1-w MP2RAGE image was bias-corrected using SPM12 and was stripped off the non-brain tissue and large sinuses using the mask obtained from the second inversion image. This pre-processed T1-w MP2RAGE was supplied as input to the high-resolution *recon-all* pipeline of Freesurfer v.6.0 (https://surfer.nmr.mgh.harvard.edu/) [63]. Additionally, the MP2RAGE T1 map was supplied as an additional input (T2-w proxy) to Freesurfer for pial surface optimisation. The segmentation and surface construction were done in the native resolution and the segmentation quality in the occipital lobe was manually curated. A probabilistic retinotopic atlas was applied to the Freesurfer reconstructed data using *neuropythy* (https://github.com/noahbenson/neuropythy) [64] to obtain participant-specific V1 and V2 regions-of-interest (ROIs) (S3 Fig b). Following the automatic segmentation and reconstruction, the WM surface was extended into WM by 30% of the cortical thickness to account for any discrepancy of the GM-WM boundary when using T1-w MP2RAGE images[65]. The first inversion image of the MP2RAGE was used to check the extended WM boundaries due to its sharp WM-GM contrast. We also extended the pial boundary by the same amount into the CSF to sample the signal away from the pial boundary. Then, we generated a total of twenty-one intermediate equi-volume surfaces within the GM using Surface tools (https://github.com/kwagstyl/surface_tools) [66] (S3 Fig c).

The functional datasets were pre-processed using Advanced Normalization Tools (ANTs) v.2.3.1 (https://github.com/ANTsX/ANTs) [67,68]. First, the functional runs were subjected to affine realignment. Next, the temporal mean of the functional run and the temporal mean of the opposite phase-encoded run were used to calculate an undistorted template image and the distortion-correction warps were saved. Lastly, a transformation matrix was calculated for each functional run to the T1-w data using the visual alignment tools in ITK-SNAP v.3.6 [69] and a final rigid alignment using ANTs. All transforms were concatenated and applied to the unprocessed functional datasets in a single resampling step using a 4th degree B-spline interpolation. This workflow was particularly developed keeping in mind the need in laminar fMRI analyses [70]. It minimises resolution losses due to multiple interpolation steps while providing the high-quality registration accuracy that is required in laminar fMRI studies.

Statistical analyses of the functional data were carried out using FSL FEAT [71,72] by modelling three regressors i.e., the stimulus design convolved with the canonical haemodynamic response function (HRF) representing the BOLD signal, the alternating label-control acquisition of the ASL sequence representing the baseline perfusion-weighting and the combination of these two regressors representing the perfusion activation. Due to the disparity in the spatial spreads of the BOLD and perfusion activation (Fig 2), a mask of the overlap between the BOLD and perfusion activation cluster thresholded masks from FEAT was created. This ensured that we sampled the BOLD and perfusion signals from the same voxels.

Laminar analyses were carried out in Freesurfer by sampling the functional time-series signal from the ROIs using nearest-neighbour interpolation. No surface or intra-cortical smoothing was applied. The laminar time-courses sampled from V1 and V2 across all participants were imported into MATLAB R2016b (MathWorks, USA) for the time-series analyses. The BOLD and perfusion-weighted time-courses were obtained for each lamina by applying surround-averaging and surround-subtraction, respectively [73–75] and the event-related average time-courses were calculated. The event-related average BOLD time-course was subsequently rescaled to percent BOLD signal change relative to the pre-stimulus baseline (~10 s). The analysis of the perfusion time-series followed several steps: First, the perfusion-weighted time-series is a measure of the modulation depth (or the magnitude of the zig-zag) of the raw ASL time-course in MRI signal units (S5 Fig). It is important to note that these data are not scaled in physiological units and is representative of the perfusion SNR of the data. We then derived the following measures from perfusion-weighted time-course: absolute and relative perfusion change, and baseline perfusion. Absolute perfusion change was calculated by taking the change in the perfusion activation (i.e., by subtracting the pre-stimulus baseline) per lamina and then normalising the signal with the mean of the EPI (to account for transmit-receive biases). The absolute perfusion change, thus obtained, is in arbitrary units but proportional to the quantitative perfusion change. The absolute perfusion change can then be rescaled into physiological units, as typically done in perfusion quantification studies [37,76]. Relative perfusion change is the percentage change in the perfusion signal due to activation per depth relative to its respective baseline. Note that the relative perfusion change does not need to be divided by the mean EPI image for scaling (as it appears both in the nominator and the denominator and thus cancels out). The baseline perfusion (Fig 1) was calculated using simple subtraction of the label-control time-points during the baseline period (~0-30 s at the beginning of the run) and pre-stimulus intervals (~0-10 s before stimulus onset) of the stimulus blocks. Laminar steady-state profiles of the BOLD signal, absolute, and relative perfusion change signals were calculated by averaging the respective signals within the ~14-28 s interval following stimulus onset. The baseline perfusion laminar profile (S4 Fig) was obtained by averaging within the entire the ROIs.

**Fig 1.**
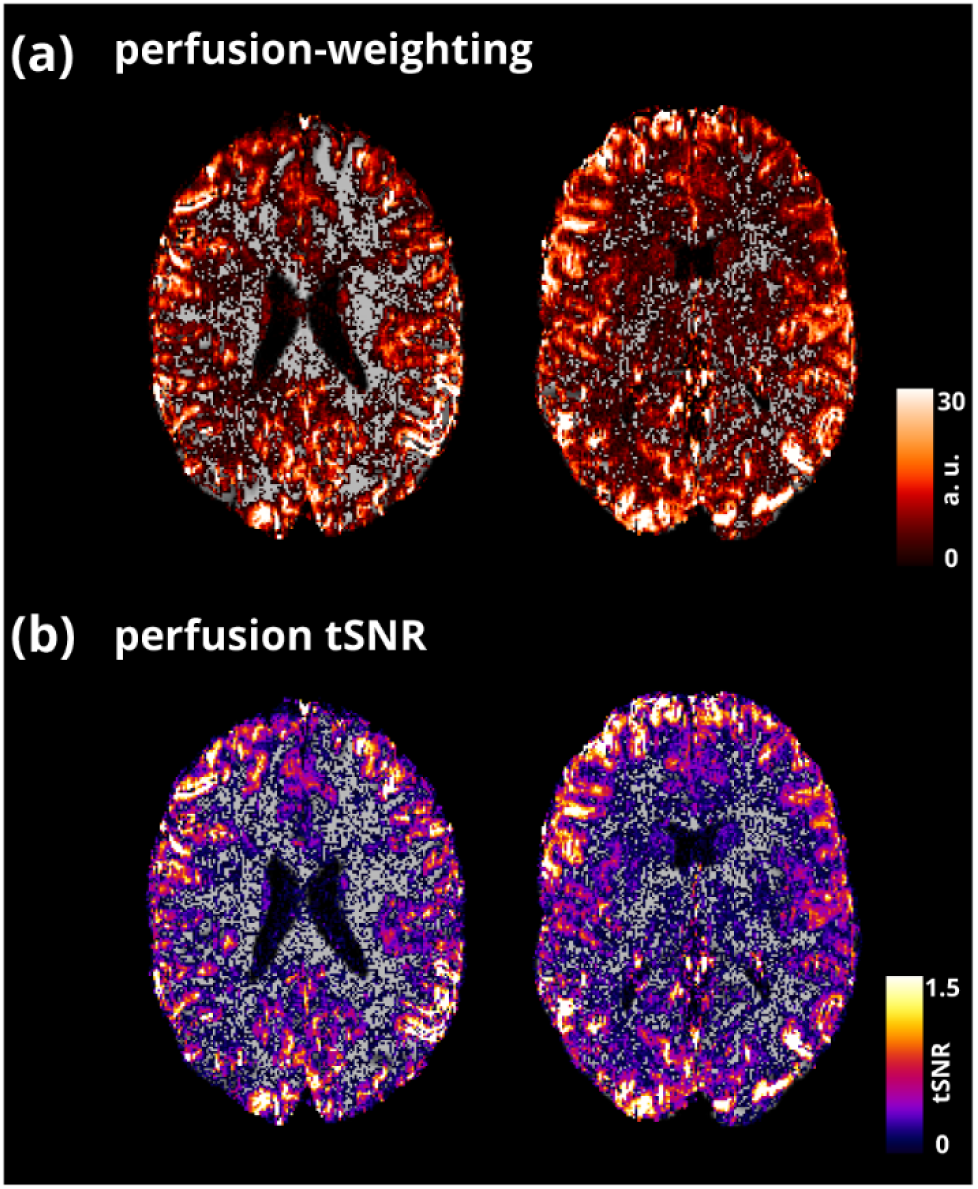
The average baseline perfusion (a) and perfusion tSNR (b) maps from a superior (left) and inferior (right) slice of an example participant is shown overlaid on the corresponding T_1_-w anatomy.

**Fig 2.**
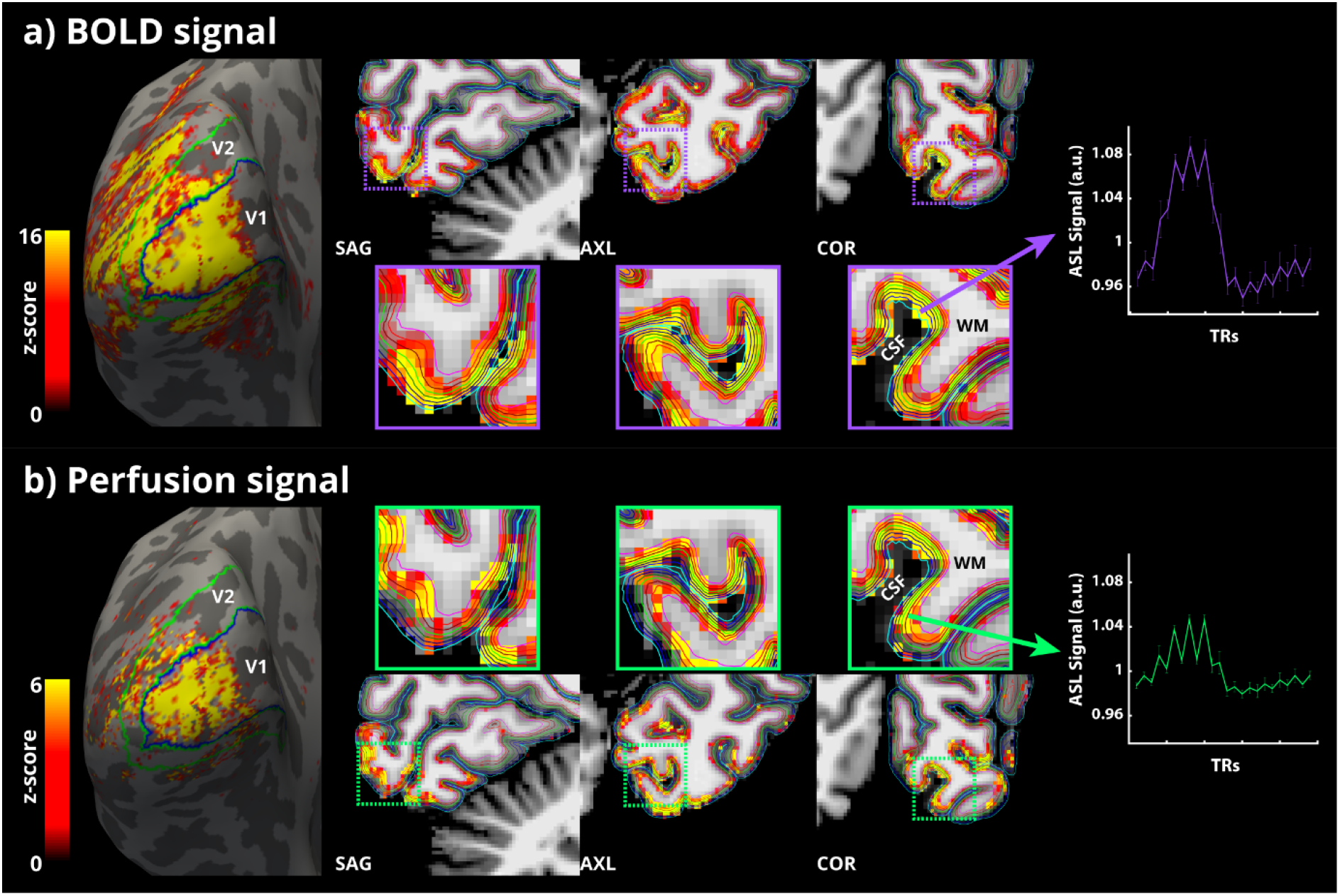
(a, left-right) BOLD signal activation map of an example participant, averaged over all runs, overlaid on the inflated left hemisphere reconstruction from Freesurfer. Contours of the V1 (blue) and V2 (green) labels obtained from Neuropythy are also overlaid on the inflated surface. Cropped orthogonal views of the participant’s occipital lobe with the BOLD signal activation map overlaid in voxel space. Boundaries of the different laminar surfaces are also overlaid, colour-coded from cyan (pial)-to-magenta (white). The purple dotted square inset indicates the zoomed-out views presented below. Event-related average ASL time-course of highly BOLD activated voxels across runs for this participant is shown to the right. (b, left-right) The same are presented for perfusion activation in green. The error-bars indicate SEM across trials.

### Simulating the laminar BOLD signal from the measured perfusion profile

The experimentally measured laminar BOLD response profiles in V1 and V2 regions were compared to theoretical predictions of the dynamical laminar BOLD signal model proposed by Havlicek and Uludağ using publicly available MATLAB code (https://github.com/martinhavlicek/laminar_BOLD_model) and plausible assumptions regarding physiology and acquisition parameters at 7T. The measured absolute perfusion laminar profiles (both baseline and activation) were used to input the physiological parameters of the model. The total amount of venous baseline CBV (CBV_0_) was set to 2 mL, of which 50% relates to microvasculature and 50% to the ascending veins. The baseline CBV (CBV_0_) distribution was set to be constant across laminae in the microvasculature but increased linearly towards the surface in the ascending veins (slope, s = 0.4). The measured baseline perfusion signal in this analysis (CBF_0_) is proportional to CBF in physiological units. Assuming that the CBV_0_ in microvasculature (1 mL /100 g) is divided uniformly between all laminae and the mean transit time through microvasculature (averaged across all laminae) is ~1 s (also see Table 2), a scaling factor *x = 5520* was estimated such that an average CBF_0_ ~60 mL /min/100 g was obtained. Given the variation of CBF_0_ across laminae, the lamina-specific transit times through microvasculature varied between ~0.6 s near the superficial laminae to ~1.2 s in the deeper laminae (0.5 s variation in transit time through capillaries between superficial and deeper laminae). The lamina-specific transit times through ascending veins were then calculated using the central volume principle (for details, see [13]). Lamina-specific changes in relative CMRO_2_ were obtained by assuming a linear coupling (n = 3) between CBF and CMRO_2_ [77]. All other parameters were defined as in the default scenario described in [13]. Please note that we did not fit the model to data but used experimentally obtained perfusion-weighted signal data and plausible biophysical parameters to generate a prediction of the laminar BOLD signal profile.

## RESULTS

### Baseline perfusion

Fig 1a and b show two representative slices (one superior, one inferior) of the average baseline perfusion map and the perfusion temporal signal-to-noise (tSNR) of a participant overlaid on the T_1_-w anatomical image. These maps show that the average perfusion signal is highly localised to the GM ribbon and demonstrates the quality of the co-registration between the acquisition slab with the anatomy as indicated by the absence of signal shifted into the ventricles and the clearly defined sulci (wherever resolvable). The perfusion-weighted data shown in Fig 1a is in arbitrary MRI signal units.

### Functional activation

Robust statistical activation was obtained for all participants for both the BOLD (Fig 2a) and perfusion signals (Fig 2b). The BOLD activation envelopes a much larger swath of cortex than perfusion activation does (Fig 2a, 2b, left panel). This is expected given the differences in the detection sensitivity (i.e. functional contrast-to-noise (fCNR)) between the BOLD and perfusion signals and the presence of BOLD signal in pial veins.

In addition, the BOLD activation obtained follows the characteristic localisation pattern observed with standard GE-EPI studies. That is, the largest BOLD activated voxels are always localised at the CSF-GM boundary (Fig 2a, purple zoomed-out boxes). In contrast, the perfusion activation was observed to be more spatially localised to the GM ribbon with the highest activated voxels localised mid- to deep-GM (Fig 2b, green zoomed-out boxes). The activation maps are shown in the three orthogonal views to highlight the consistency of the GM localisation of activation in 3D.

Finally, the ASL time-courses exhibit a zig-zag modulation that is characteristic of ASL sequences (due to the acquisition of alternating label and control volumes) demonstrating the high quality of the data. The modulation depth of this zig-zag represents the amount of labelled spins delivered to the tissue and is, therefore, proportional to tissue perfusion. In Fig 2 (purple), the ASL time-course obtained from the highest BOLD signal activated voxels shows the typically observed increase in the BOLD signal magnitude during activation with weaker zig-zag modulation. On the other hand, the ASL time-course obtained from the highest perfusion-activated voxels (Fig 2, green) shows the strong zig-zag modulation throughout but with lower BOLD signal modulation. All three key differences between the BOLD and perfusion activation signals were consistently observed in all the participants.

### Laminar analysis

The group-average laminar time-courses of BOLD signal change, absolute perfusion change, and relative perfusion change are shown in Fig 3 for V1 and in Fig 4 for V2. The temporal behaviour of the three sampled signals across all laminae is presented as a heatmap in the top row with time along the X-axis and the cortical depth along the Y-axis and the magnitude of the signal in colour code. We observed inter-regional differences with laminar responses of all three signals, with V2 having a lower amplitude than V1. The laminar profiles of the BOLD signal change exhibit positive slopes (Slope V1: 4.88 ± 0.129, Slope V2: 4.81 ± 0.195) with a strong linear trend (*R^2^* V1: 0.986, *R^2^* V2: 0.967). The laminar profiles of the relative perfusion change, on the other hand, exhibit negative slopes (Slope V1: −4.91 ± 0.27, Slope V2: −4.64 ± 0.111) with a strong linear trend (*R^2^* V1: 0.939, *R^2^* V2: 0.988). Interestingly, the absolute perfusion changes exhibit a moderately positive slope (Slope V1: 3.35 ± 0.68, Slope V2: 3.80 ± 0.485) albeit without a strong linear trend (*R^2^* V1: 0.537, *R^2^* V2: 0.745). In the perfusion signals, slight oscillatory behaviour is observed during the post-stimulus period.

**Fig 3.**
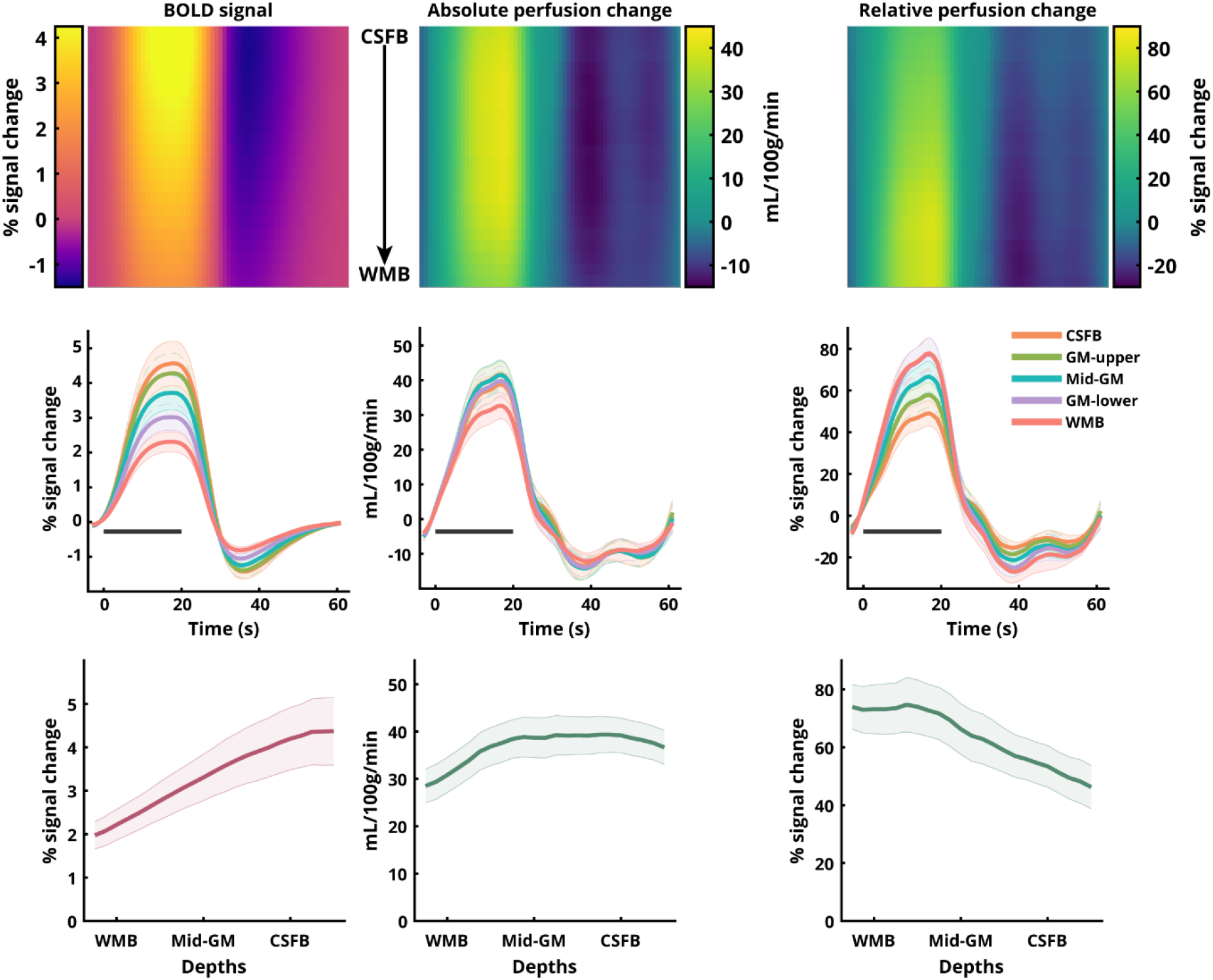
Laminar BOLD and perfusion signal changes in human V1 ROI obtained using sub-millimetre 3D-EPI ASL at 7T. (top row) Heatmap representations of the group-average BOLD signal change, absolute perfusion change, and relative perfusion change with cortical depth along Y-axis and time along the X-axis. (middle row) Five out of the twenty-three total laminar time-courses for the respective sampled signals. (bottom row) Laminar profiles of the positive responses for the respective sampled signals with cortical depth along X-axis. Error-bars indicate SEM across participants. The black bar in the middle row indicates the stimulus duration.

**Fig 4.**
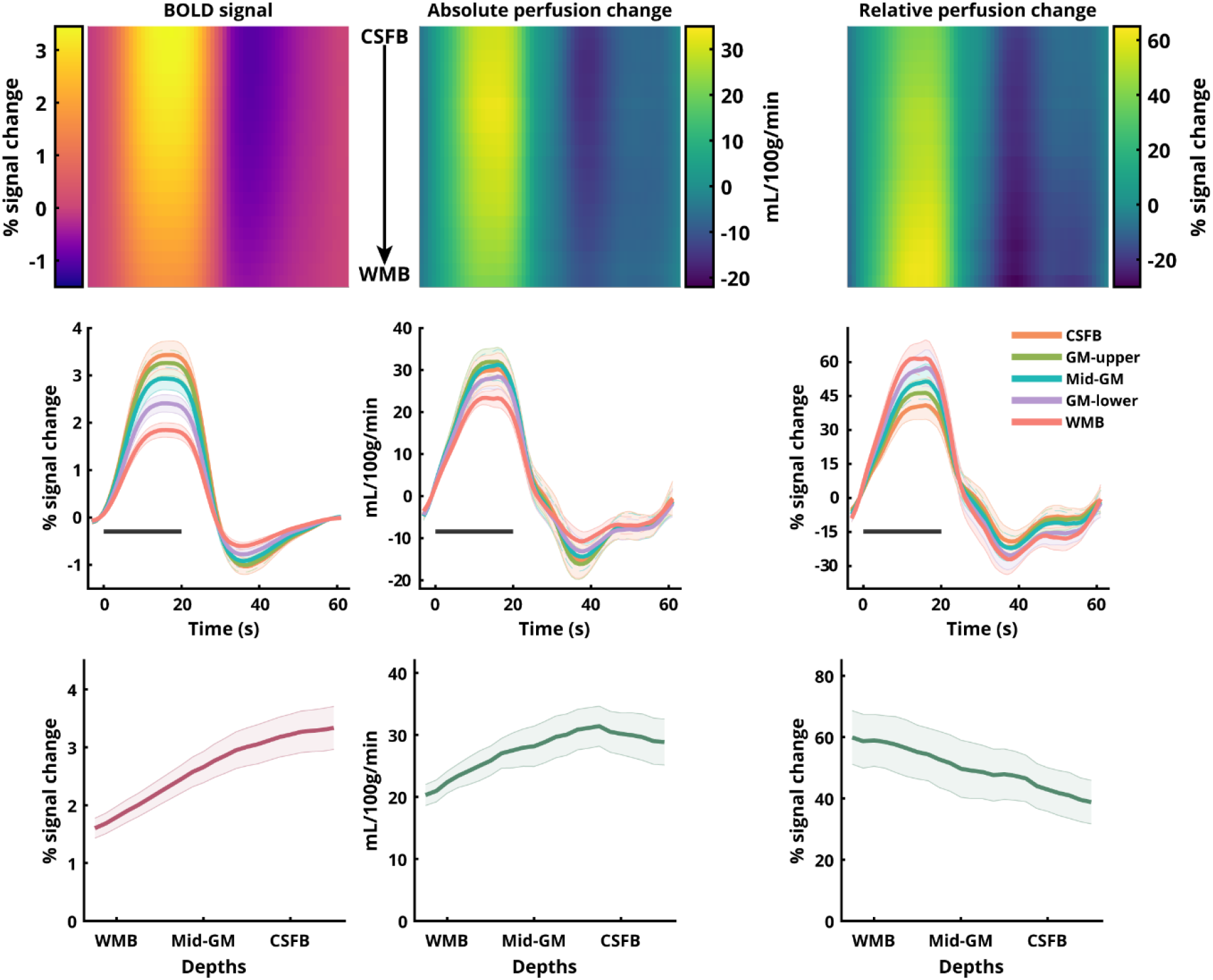
same as Fig 3 for V2 ROI.

### Simulations of the laminar BOLD signal

Fig 5 shows the simulated laminar BOLD signal profile (solid blue lines) and the experimentally measured laminar BOLD signal profiles (dotted purple lines, see Fig 3, 4). The measured and simulated profiles were highly congruent (Pearson’s correlation: r = 0.9984 for V1, r = 0.9977 for V2), demonstrating that, despite the discrepancy of the relative & absolute perfusion and the BOLD signal profiles, they are in fact compatible with each other.

**Fig 5.**
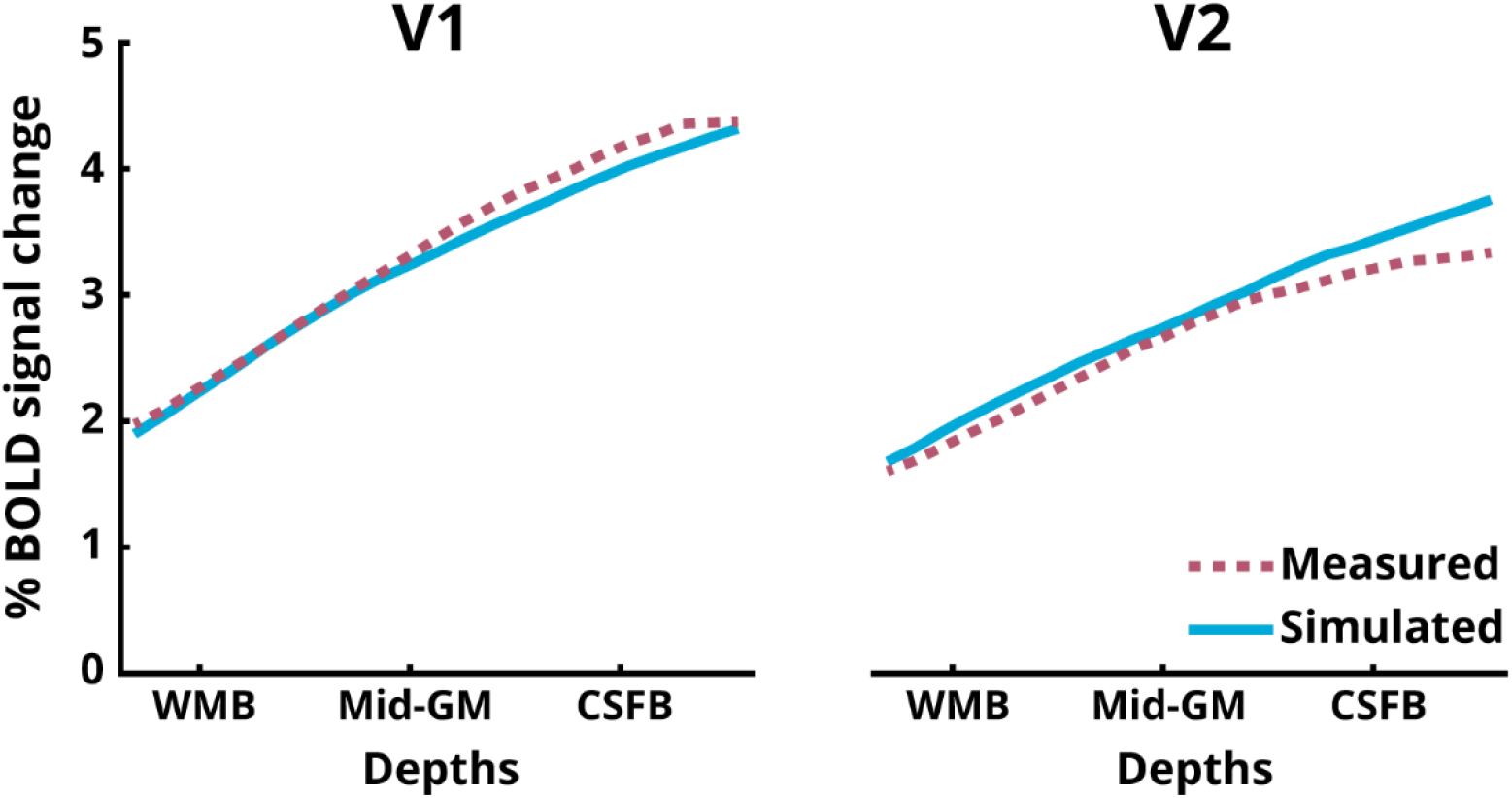
Laminar profiles of the BOLD signal for V1 and V2 ROIs. The measured responses are the same as the BOLD signal profiles in Fig 3 and 4. The simulated profiles are obtained using the dynamical laminar BOLD model [13].

## DISCUSSION

Here, we demonstrated, for the first time in humans, isotropic sub-millimetre spatial resolution perfusion fMRI using ASL. We found incongruent cortical depth profiles between the BOLD signal and perfusion changes, which, however, turned out to be physiologically consistent with each other after employing a dynamical BOLD signal model.

### Functional BOLD and perfusion activation

We obtained robust participant-specific, single-session activation maps for simultaneously acquired isotropic sub-millimetre spatial resolution BOLD and perfusion signals at 7T. We observed a larger spread of activation for the BOLD signal (Fig 2a) compared to the perfusion signal (Fig 2b). This is expected because the detection sensitivity of the perfusion signal (fCNR) is much lower than that of the BOLD signal [33,37,38]. Additionally, this can also be explained by the higher spatial specificity of the perfusion signal compared to the BOLD signal, which is susceptible to non-local signal spread due to downstream venous bias away from the actual site of activation [14]. This is also observed in high-resolution fMRI with the highest BOLD activated voxels located at the CSF-GM boundary (Fig 2a). On the other hand, the perfusion activation map exhibits a well-defined localisation to the cortical ribbon (Fig 2b), mostly located in cortical GM [18]. Importantly, given that perfusion signal has much lower fCNR than the BOLD signal in standard resolution studies (2-4 mm in each direction), it was not necessarily expected that ASL will have enough sensitivity at submillimetre resolution for detecting perfusion activation. One reason that with increasing resolution there is enough perfusion fCNR is that not only image SNR but also partial voluming with CSF and WM is decreased, i.e. thermal and physiological noise coming from outside GM are reduced. This is different for the BOLD signal as pial vessels located in CSF (see Fig 2a) do contribute to the overall BOLD signal in low resolution studies and therefore increases in spatial resolution decrease both image SNR and overall signal contribution. That is, going from low- to high-spatial resolution penalizes CNR of the BOLD signal more than of the perfusion signal.

Recently, a novel fMRI approach called VAPER [78] has also been put forward as a contrast useful for perfusion-weighted high-resolution fMRI by mixing VASO and perfusion contrasts. Although the combination of two contrasts boosts VAPER’s sensitivity, it markedly complicates its ability to quantify perfusion but also, its physiological specificity. Thus, established ASL techniques remain the most feasible way to acquire in *vivo* perfusion-weighted images that can be straightforwardly validated using quantitative fMRI models, and can be expected to provide reproducible results across a wide range of sequence parameters and field strengths [33]. In this regard, the present study is the first demonstration in humans of the improved spatial specificity of the perfusion signal compared to the BOLD signal using ASL at a sub-millimetre spatial resolution.

A recent review of non-BOLD laminar fMRI methods illustrated the potential of the 3D-EPI PASL sequence for perfusion-weighted laminar fMRI applications at ultra-high field. As highlighted in the review, the perfusion contrast has been highly desired for laminar fMRI [79] as the perfusion signal is relatively unaffected by the venous compartments, both by the pial and ascending veins, and the large arterial compartments. In comparison, the BOLD signal is heavily weighted towards the venous compartments and the VASO signal can have contributions from both arterial and venous in addition to microvasculature CBV changes [34,80,81]. The reason for the high perfusion localisation specificity is that the tagged arterial water is mostly exchanged with the tissue at the level of the capillaries. In addition, the transit delay for the labelled blood to arrive at the region-of-interest (in this case, occipital lobe) can be ~1-1.3 s [82]. Together with the blood transit time within tissue on the order of ~1-2.5 s [83] only little longitudinal magnetisation of the tag remains (due to T1 decay), i.e. that almost no magnetisation of the label is present in venous blood (see S8 Fig), except for artefacts caused by labelling of venous blood superior to the imaging slab in some ASL schemes (see [76]). The transit time for the acquisition in the present study was optimised for the visual cortex and is reflected in the inter-regional differences in the baseline perfusion signal and its temporal stability of the tissue (Fig 1). The absence of the venous bias and the signal being dominated by the capillary compartment implies that the perfusion contrast more closely follows both the spatial profile and the amplitude of cortical metabolism and neuronal activation. Another important aspect of ASL acquisitions is the possibility to obtain a quantitative estimate of the baseline signal across depths.

The difference between the highly BOLD-activated or highly perfusion-activated voxels is readily visible in the ASL time-courses (see Fig 2). The time-courses for perfusion activation show reduced amplitude of the signal envelope and larger difference between pairs of data points (i.e. the zig-zag modulation) indicating that these voxels contain signals from mostly the microvasculature and that observed responses are indeed capturing the changes in perfusion. In contrast, there are small zig-zag changes relative to the overall signal envelope in the time-courses for the highest BOLD activation, reflecting a smaller contribution from microvasculature. This means that the spatial non-overlap that we observe between the perfusion and BOLD signals is driven largely by differences in the underlying physiology and not the differences in SNR.

### Laminar BOLD and perfusion responses

We replicated previous findings [21,84,85] that the event-related average BOLD signal amplitude (Fig 3 and 4, first column) increases towards the CSF-GM boundary (e.g., [86,87]). The BOLD signal increase to the superficial layers is well understood and can be attributed to two signal biases: a) increase in baseline CBV of the intra-cortical ascending veins and b) the non-local blooming effect from the pial veins ([88], and for overview see, [2]). The presence of these biases in the BOLD signal makes the interpretation of the measured laminar signal profile, specifically in the superficial layers, challenging [89]. One approach to deal with the issue of spatial bias in GE-BOLD signal is model-driven spatial “deconvolution” [22,24], which, however, has not yet been validated with another (simultaneously) acquired fMRI modality.

The profile of the relative perfusion change (Fig 3 and 4, right column) exhibits the opposite behaviour (compared to the BOLD profile) with the magnitude of the signal increasing towards the GM-WM boundary with a strong linear trend. Furthermore, QUIPSS II pulses were employed in the present study allowing clear-cut definition of the tagged bolus. This means that the observed patterns of laminar signal behaviour are unlikely to be due to undelivered tagged blood in the diving arterioles. Although the impact of the QUIPSS II pulse depends on the chosen parameters and the arrival times to the regions-of-interest, an increase in blood flow upon activation can result in a more complete delivery of the tagged spins to the tissue, including the deeper layers at the time of volume acquisition. This could yield a larger fractional perfusion change in the deeper layers relative to the baseline condition. While it is usually argued that for feed-forward stimuli the peak in activation must be in the middle layers, electrophysiological evidence, histology, and a previous BOLD signal study after spatial “deconvolution” (Marquardt et al., 2018) support the view that V1 also receives high input into layer VI in addition to layer IV. Please note that despite the high spatial resolution used in this study, we do not detect a fine-grained distinction between laminae. The perfusion spatial profile obtained, thus, represents a smoothed version of the underlying neuronal activity. For example, data shown in Fig 4b and 4d in [22] and in [90] (see Fig 9 in [22]) are compatible with the spatial profiles found in the current study. We find that the relative increase in the perfusion signal in the middle to deeper layers is also consistent with animal literature (see also [18,91,92].

The absolute perfusion signal change profile (Fig 3 and 4, middle column) exhibits a weak positive slope and non-linear behaviour across depths. However, both relative and absolute signal changes are derived from the same perfusion-weighted signal obtained after surround-subtraction and the difference stems from the spatial profile of the baseline perfusion (S5 Fig). Please note, that the increase of the absolute perfusion signal from WMB to CSFB (by ~30-50%) is much smaller than that of the BOLD signal (by ~100-120%). Additionally, in contrast to the BOLD signal, the absolute perfusion change drops beyond the CSF border. Taken together, the relative and absolute perfusion signal changes differ in their depth-dependent behaviour and both differ from the BOLD signal either in the sign of their slope or the relative increase of the profile towards the surface.

In order to test if this discrepancy between the relative & absolute perfusion profiles and BOLD profile can be reconciled, we simulated the BOLD signal profile from the measured perfusion profiles using the recent dynamical laminar BOLD signal model (for details, see S4 Fig). We show that the positive slope and the relative increase of the measured BOLD profile can be obtained from the laminar profile of the relative (having negative slope) and absolute (having much smaller increase towards the surface) perfusion signal by modelling the ascending vein bias, i.e., simulating the laminar BOLD response in a forward manner. Therefore, we conclude that despite their seemingly contrasting behaviours, the BOLD and perfusion signal profiles are, in fact, physiologically consistent with each other.

Additionally, the BOLD time-courses exhibit a strong post-stimulus undershoot (PSU) consistent with previous studies [84,93]. Interestingly, our perfusion measures also exhibit PSUs but with smaller amplitudes relative to the positive response. In contrast to the smooth recovery of the PSU to baseline in the BOLD signal, the perfusion PSU exhibits slight oscillatory behaviour. These post-stimulus oscillatory transients are consistent with previous reports of perfusion measurements in humans (e.g., [94]) and with optical imaging in rodents [95]. The oscillatory transients observed in the previous perfusion study in humans [94] could not resolve any depth-dependent modulations owing to its much lower spatial resolution (i.e. 2.65 × 2.65 × 5 mm^3^). The post-stimulus oscillations in our study near the WM boundary are smoother and evolve with a different oscillatory phase than near the CSF boundary, where the oscillations are more pronounced (Fig 3 and 4, middle panels). While it is interesting, pin-pointing the exact vascular physiology that elicits this behaviour is beyond the scope of this study.

Taken together, we believe that the current study presents a breakthrough in non-BOLD fMRI research with the development of sub-millimetre resolution perfusion fMRI using ASL and its feasibility for layer-specific investigations, which has hitherto been an uncharted territory in humans.

### Data processing

We developed a novel workflow to pre-process anatomical images (S1 Fig) by using the second inversion image of the MP2RAGE and SPM’s segmentation to automatically mask out the sagittal and transverse sinuses that are crucial for highly accurate pial surface delineation using Freesurfer’s recon-all. In some participants, the workflow required (albeit very little) manual corrections of the segmentation masks. Additionally, we supplied the quantitative T1 map of the MP2RAGE as a proxy T2 in the recon-all stage 3 to further improve the pial surface reconstruction. We used an open-source python package *neuropythy* [64] to apply a probabilistic atlas of retinotopy in participant’s native space to generate automatic labels of V1 and V2 (S3 Fig b). We qualitatively compared this atlas-based approach on a separate dataset of pRF mapping that was acquired using the same scanner, head coil and similar coverage (data not shown) and found high degree of overlap. Please note, the focus of the present work is distinguishing the BOLD and perfusion signals and does not rely on the perfect delineation of V1 and V2 borders. Cortical layering was done using the equi-volume approach [96] using *Surface tools* [97] as equi-volumetric layering is not natively supported in Freesurfer. Nevertheless, for spatial resolutions such as the present study the exact choice of the layering model does not affect our main conclusions [98]. Even though the layering is done on the whole cortical ribbon, we manually ensured that the delineations were accurate within the V1 and V2 masks in each participant. As ASL measures the BOLD signal simultaneously with perfusion, the BOLD signal profile serves as an internal control for the accuracy of the segmentation and layering. The BOLD signal spatial profile for feed-forward stimuli (such as checkerboards as used in this study) is well known and the BOLD signal derived from the ASL data reproduces this well-known amplitude increase towards the surface of the cortex (see Fig 3 and 4), confirming the accuracy of the data processing in this study.

We have previously encouraged studies to align anatomical-to-functional data in order to reduce the blurring due to the application of several resampling steps on the high-resolution, high-fidelity laminar fMRI datasets [84]. While minimal processing approaches for fMRI have been proposed [99], as far as we know, they have not been applied to laminar fMRI. To this end, our workflow using the ANTs framework estimates, combines and applies transformations of motion, distortion-correction and co-registration to the anatomical image in a single resampling step, thereby reducing the amount of smoothing resulting from the processing of the data. Please note, in some ASL acquisitions there may be strong differences in image contrast between the label and control images and the choice of realignment cost-function may impact the quality of correction. However, this was not the case for the present study (S7 Fig). Importantly, as can be seen in Fig 2, high values of perfusion were tightly confined to the GM ribbon illustrating the high accuracy of the segmentation and co-registration in the present study.

### Limitations

The goal of the present study was to demonstrate the feasibility of using perfusion-weighted contrast with ASL for laminar fMRI and, to that end, we employed a block design with a strong feed-forward visual stimulus that is known to elicit widespread activation. Due to the lower SNR of the perfusion-signal, we averaged 40 min worth of functional runs. While there is the undoubted benefit in spatial specificity, ASL may not be well-suited for all laminar fMRI studies, particularly those with small effect sizes. In addition, GE-BOLD laminar fMRI data are routinely acquired with 0.6-0.8 mm isotropic resolutions, higher than the current ASL study. While the use of Partial-Fourier acquisition can reduce the effective spatial resolution along a dimension, the amount of blurring was reduced by using POCS reconstruction algorithm instead of zero-filling. Nevertheless, to the best of our knowledge, this study remains the highest spatial resolution functional ASL study in humans till date. Going forward, sub-millimetre resolution ASL can be invaluable to studies that are examining BOLD signal physiology, for validating existing models or for brain areas contaminated by close large pial veins.

The lowest achievable TE in the present study was 15 ms owing to the EPI readout, which is not ideally suited for perfusion imaging. Although, it would be desirable to achieve shorter TEs (e.g. ~3 ms or less) for better perfusion-weighting, it is currently not possible using conventional Cartesian EPI. To this end, there has been recent progress in non-Cartesian (e.g. spiral readouts) ASL fMRI at ultra-high field [100,101]. Dual-echo spiral acquisitions can be particularly useful for simultaneous perfusion and BOLD imaging achieving the first echo at ~2 ms (perfusion-weighted) and the second echo at ~25 ms (BOLD) at 7T. However, these non-Cartesian acquisitions are prone to inaccuracies in the spiral trajectories due to gradient imperfections that require real-time monitoring and correction using specialised field-monitoring hardware [100]. However, research and development are still underway to address these technical challenges in non-Cartesian imaging and currently sub-millimetre fMRI acquisitions have not been demonstrated.

We show that high-resolution ASL at ultra-high field is possible using the standard commercial head-coil with single-channel transmit (NOVA Medical, USA). However, the B_1_^+^ inhomogeneities remain a major hurdle [33]. While We were able to mitigate this to some extent using dielectric pads [50,51], Future studies will be able to take advantage of advances in parallel transmission (pTx) technology [102] or the use of dedicated labelling-only RF-coils [103–106] to potentially further optimise high-resolution ASL fMRI at ultra-high fields. Having demonstrated the feasibility of perfusion-weighted laminar fMRI using ASL at a sub-millimetre spatial resolution, future studies will be able to systematically evaluate different properties ASL and its impact on the perfusion signal evolution at ultra-high field.

## ACKNOWLEDGEMENTS

The study was supported by the Netherlands Organisation for Scientific Research (NWO) VIDI grant (452-11-002) and the Institute for Basic Science, Suwon, Republic of Korea (IBS-R015-D1) to K.U., NWO VENI grant (016-198-032) to L.H., NWO VIDI grant (016-178-052) to and National Institutes of Health grant (R01MH111444, PI: David A. Feinberg) to B.A.P. The funders had no role in study design, data collection and analysis, decision to publish, or preparation of the manuscript. All data were acquired at the Scannexus B.V. facilities, Maastricht, The Netherlands.

## DATA AND CODE AVAILABILITY STATEMENT

The authors do not have permission to share the data. The 3D-EPI PASL sequence used to acquire the data is available upon request via a SIEMENS C2P agreement. All code used for analysis are either already publicly available or will be made available upon publication.

