## Supplementary Figures for "Sub-Millimetre Resolution Laminar Fmri Using Arterial Spin Labelling in Humans at 7 T"

### SUPPORTING FIGURES

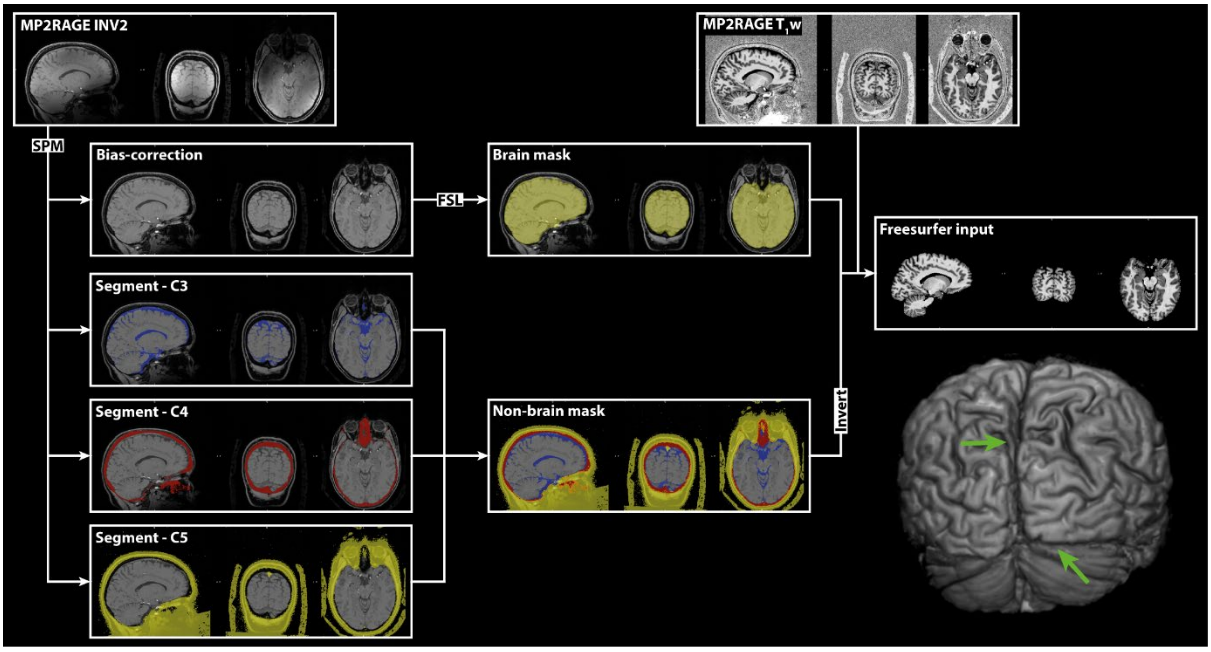


S1 Fig. Anatomical data pre-processing workflow for MP2RAGE data for input to Freesurfer's recon-all. Tissue classes C3-C5 were thresholded on a subject-by-subject basis to include as much of the sagittal and transverse sinuses as possible. In most of our subjects, this procedure did not require any manual correction of the combined mask. ITK-SNAP v. 3.6 (Yushkevich et al., 2006) was used to make any manual corrections when required. The Freesurfer T1-w input is presented as a 3D render showing an intact GM surface at the occipital lobe with the green arrows indicating locations of the now automatically stripped sagittal and transverse sinuses.

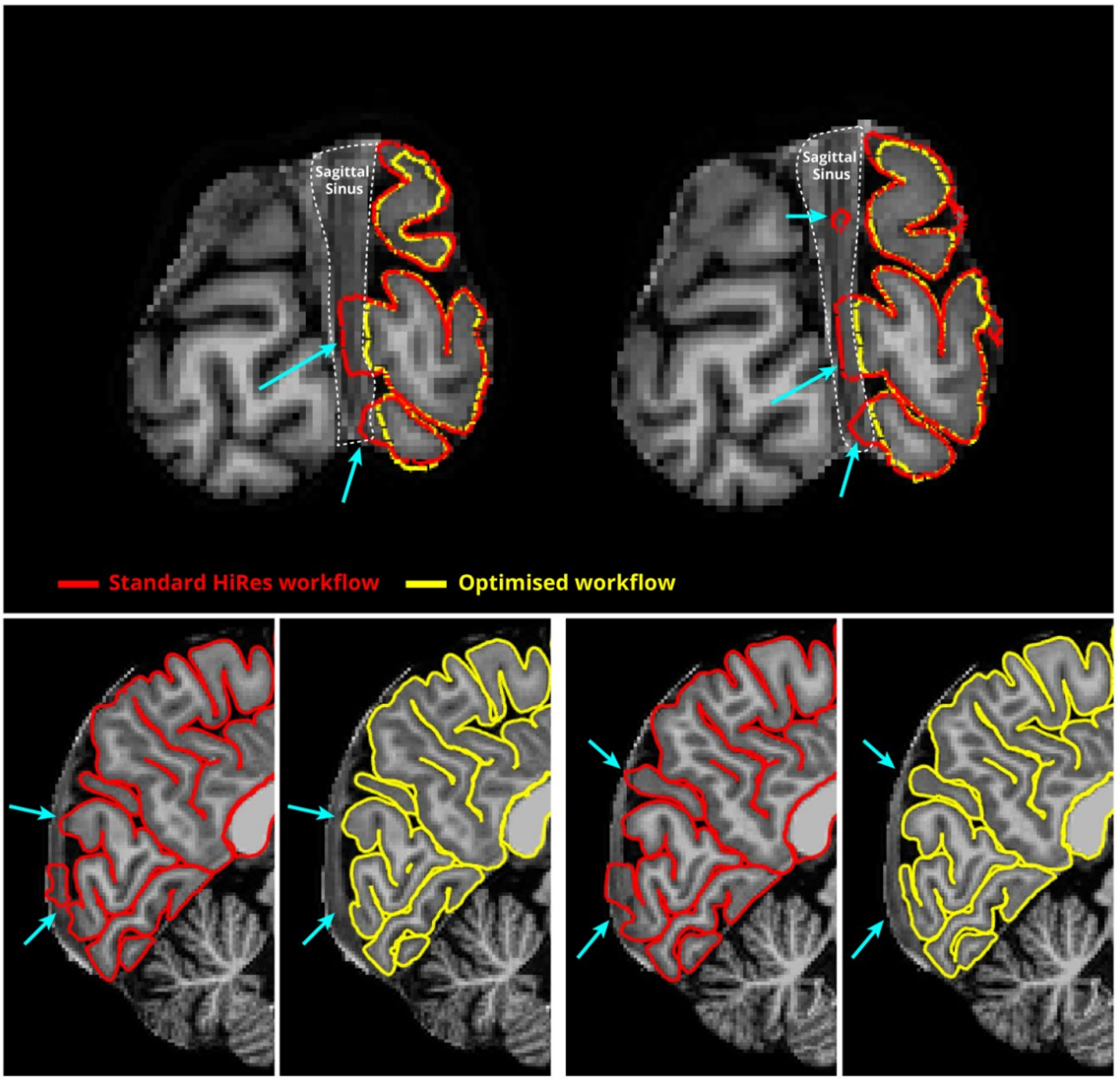


S2 Fig. (top) Coronal view of the pial surface overlaid on a standard skull-stripped T1-w image. Shown in red is determined from the standard HiRes workflow and in yellow is determined from the optimised workflow shown in Supplementary Fig 1. White dotted lines indicate sagittal sinus and cyan arrows emphasise the erroneous placement of the pial surface in the standard workflow. (bottom) Sagittal views of the pial surface from the standard and optimised workflows to further illustrate their differences in outcome indicated by the cyan arrows.

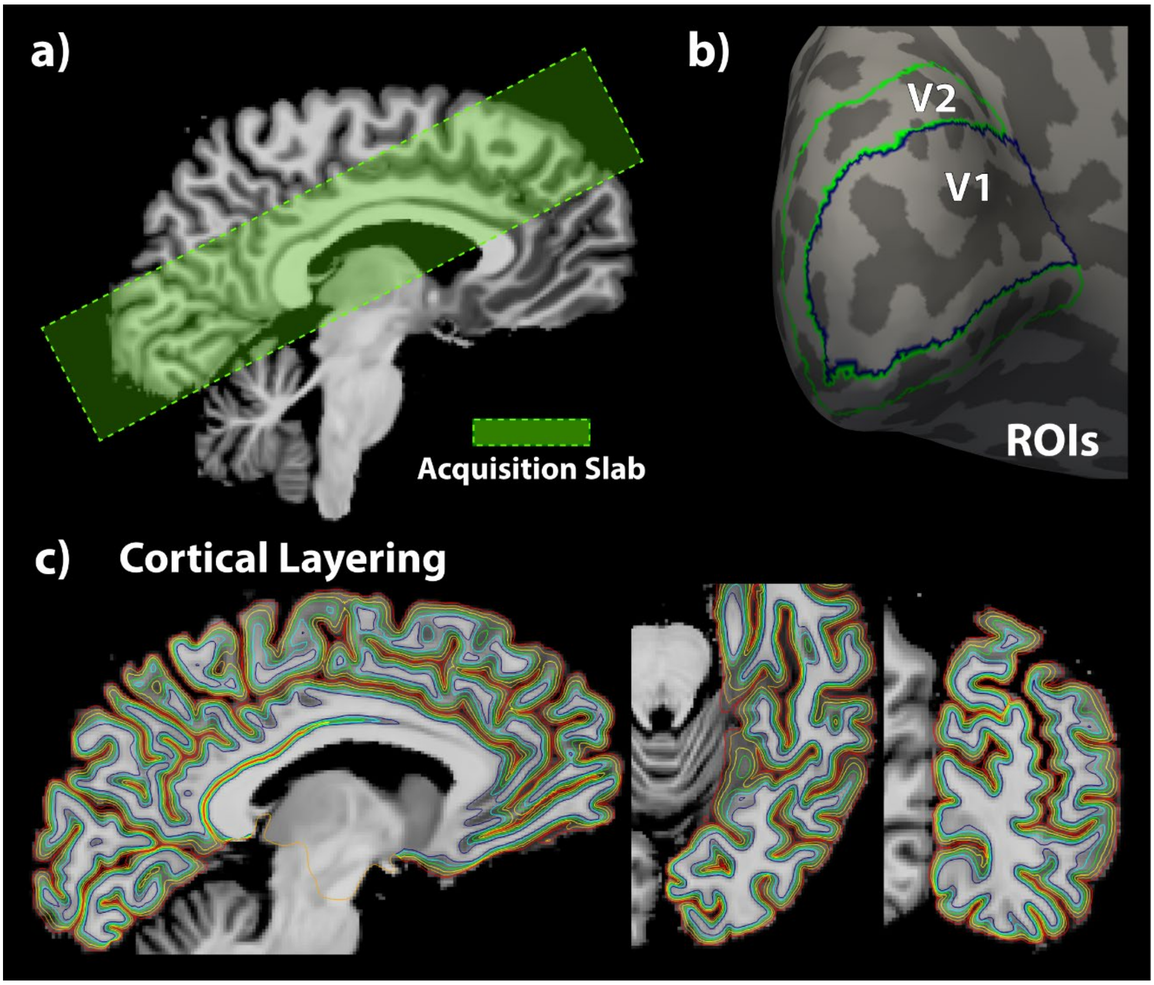


S3 Fig. Illustration of (a) the extent and orientation of the acquisition slab (b) the atlas-based delineation of the V1 and V2 ROIs (c) a subset of all laminar surfaces overlaid on one hemisphere.

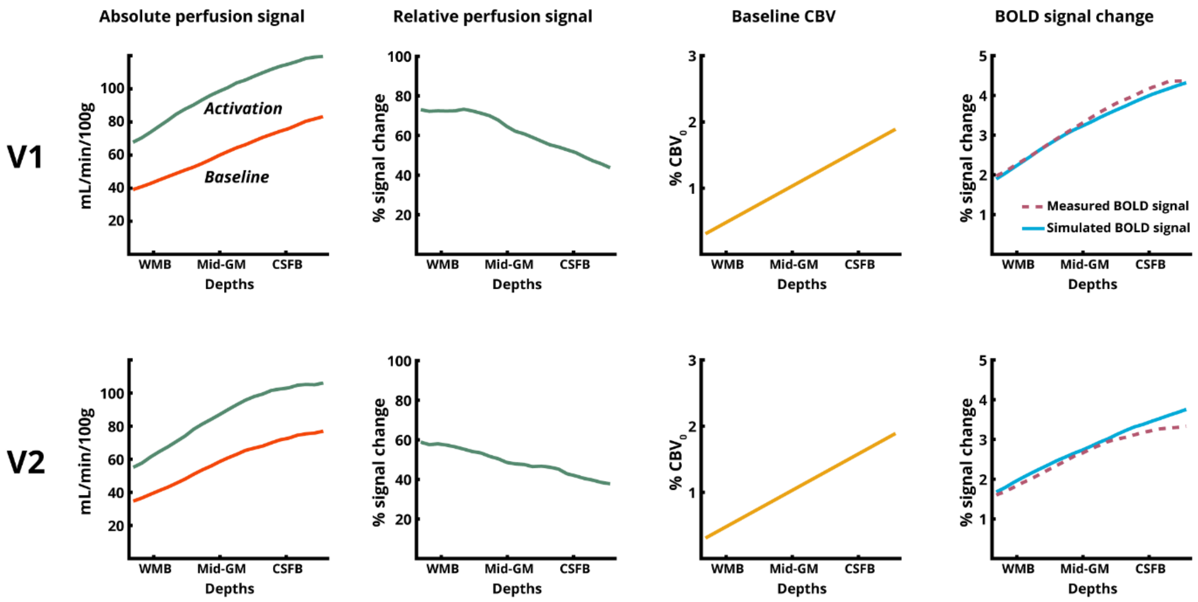


S4 Fig. (left-to-right) Measured absolute perfusion signal for activation and baseline, relative perfusion laminar profile obtained from measured absolute perfusion data, assumed laminar baseline blood volume, and comparison of measured BOLD signal change and simulated BOLD signal change as predicted by the dynamical laminar BOLD model (Havlicek and Uludağ, 2020).

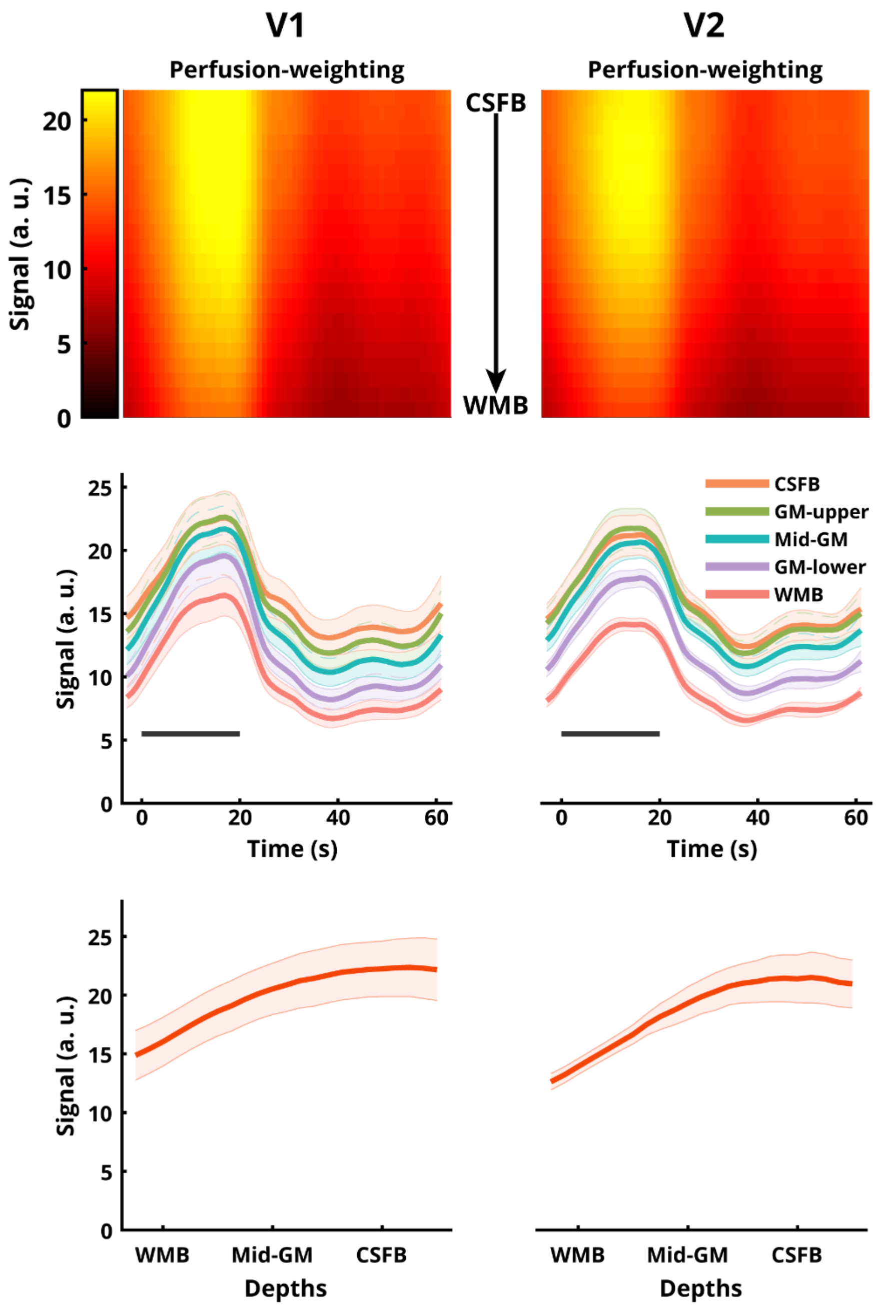


S5 Fig. Laminar profile of the perfusion-weighted signal (obtained after surround-subtraction) in human V1 and V2 from the sub-millimetre 3D-EPI ASL at 7T. (top row) Heatmap representations of the group-average perfusion-weighted signal with cortical depth along Y-axis and Time along the X-axis. (middle row) Five out of the twenty-three total laminar time-courses and (bottom row) laminar profiles of the positive response for the perfusion-weighted signal. All error-bars indicate SEM. The grey bar in the middle row indicates the stimulus duration.

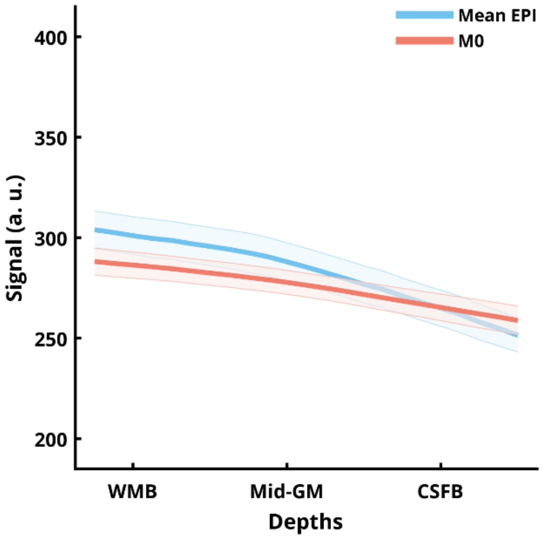


S6 Fig. Laminar profile of the mean EPI from the sub-millimetre 3D-EPI ASL and long TR calibration image (M0) at 7T.

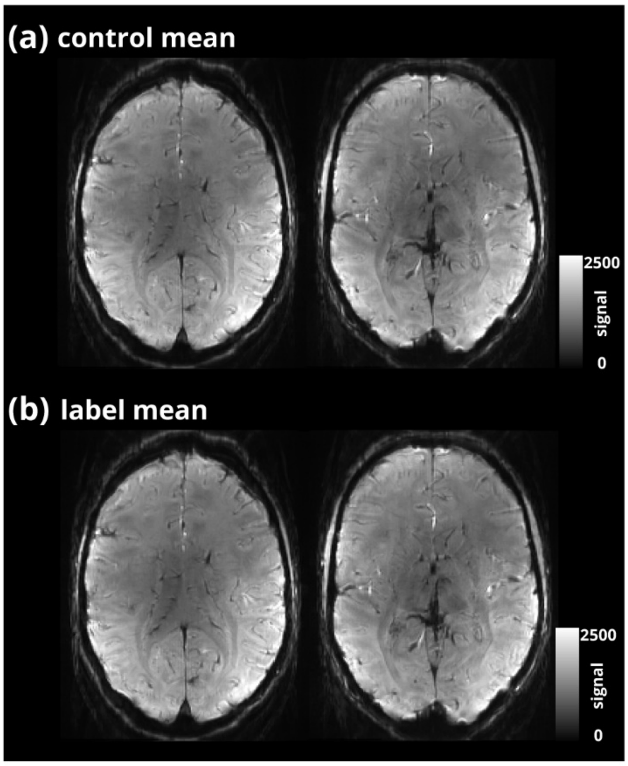


S7 Fig. Temporal mean EPI image of all control (a) and label (b) volumes from a single ASL run.

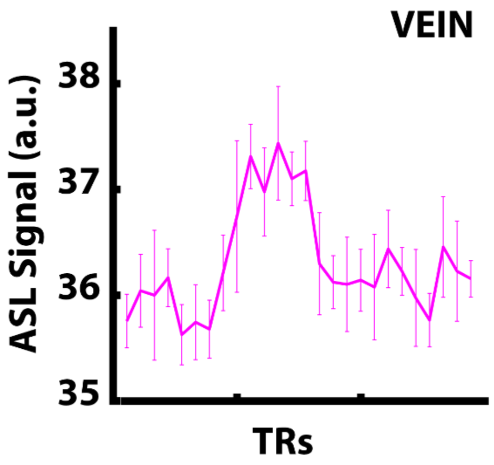


S8 Fig. Event-related average ASL time-course sampled from the sagittal sinus near V1. Error bars indicate SEM across trials. Please note the near absence of the characteristic ASL zig-zag signal modulation and the scale of the Y-axis.
